## Supplementary figures and images for "The ortholog of human REEP1-4 is required for autophagosomal enclosure of ER-phagy/nucleophagy cargos in fission yeast"

### Supplementary Figure S1

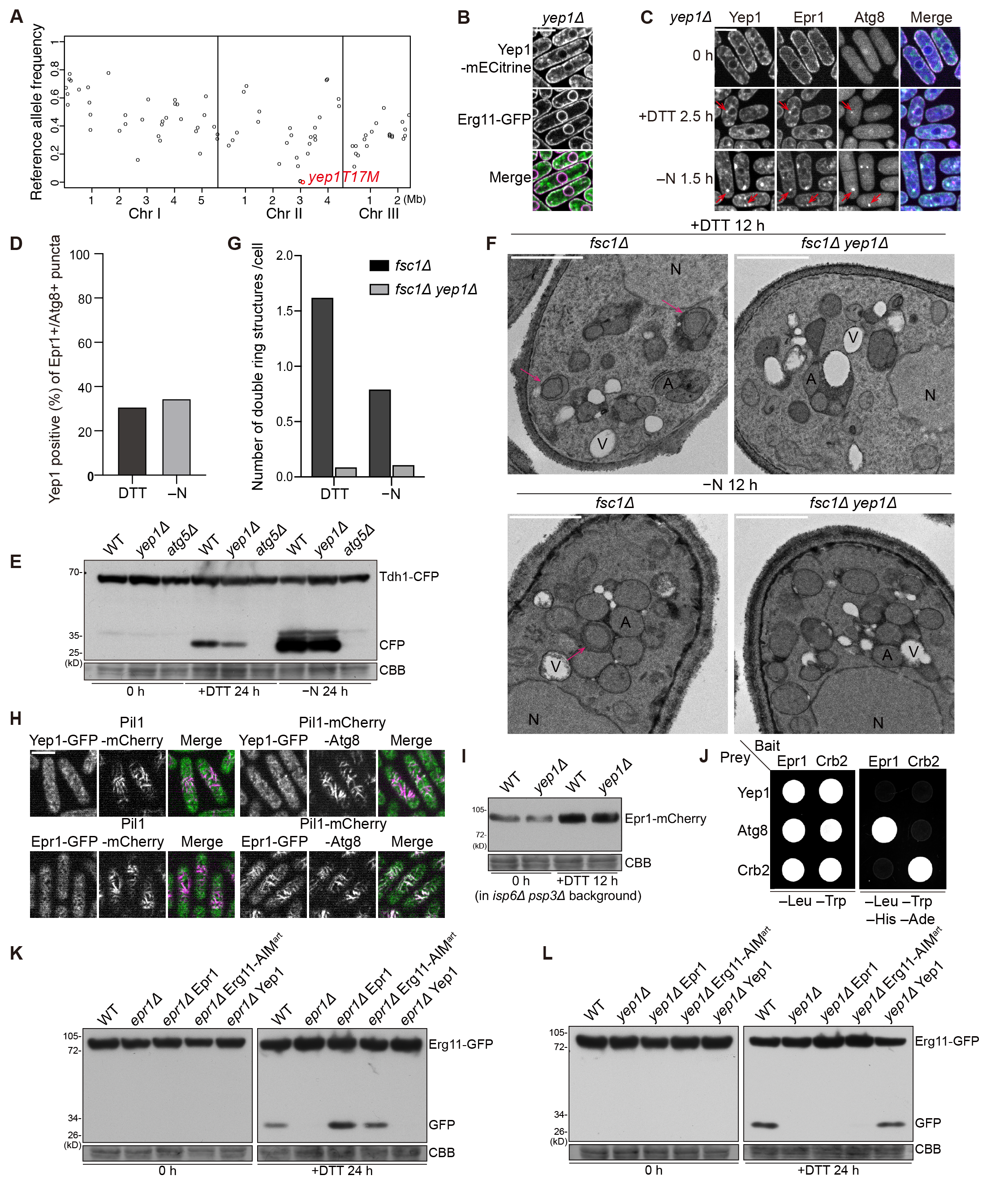

### Supplementary Figure S2

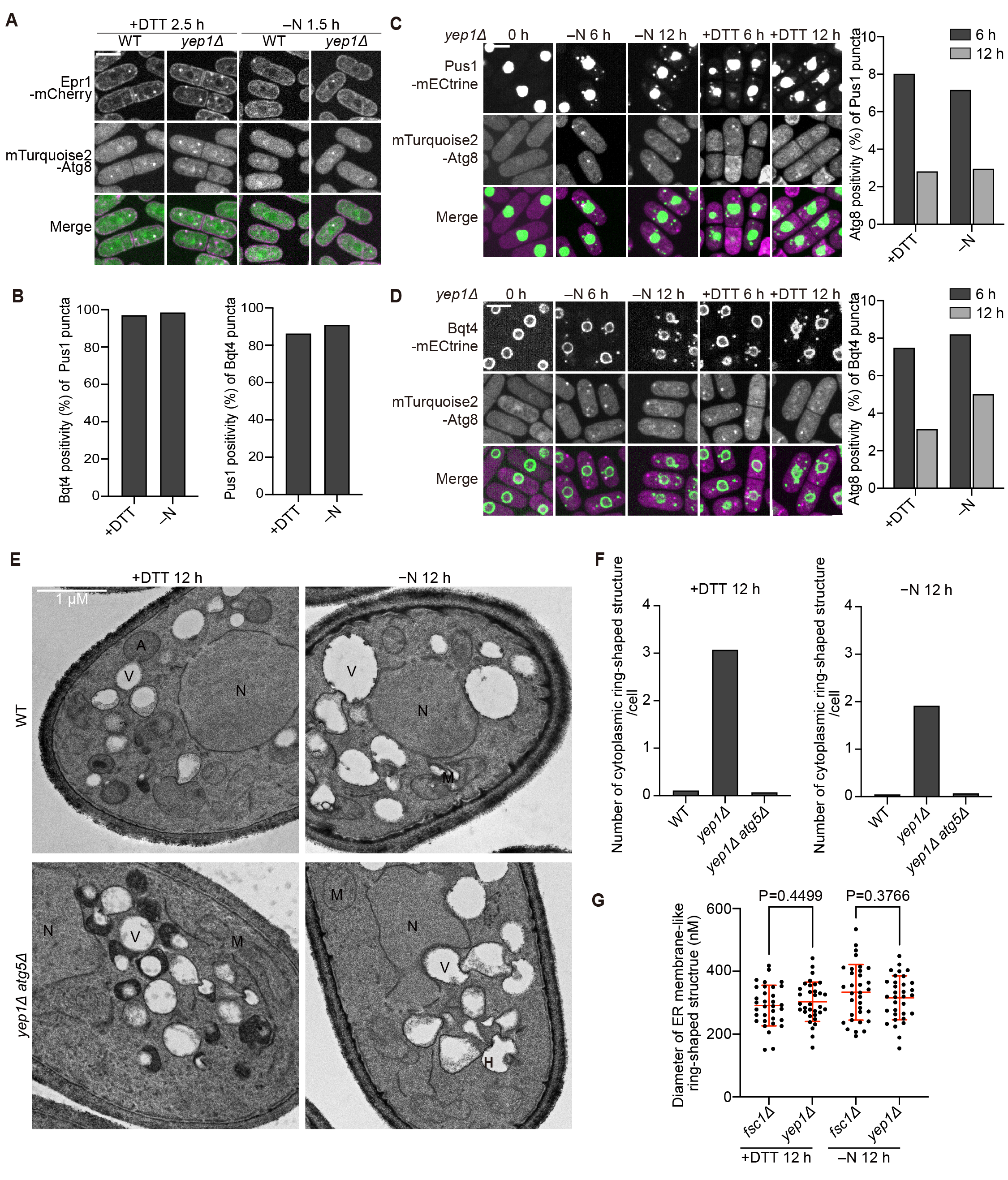

### Supplementary Figure S3

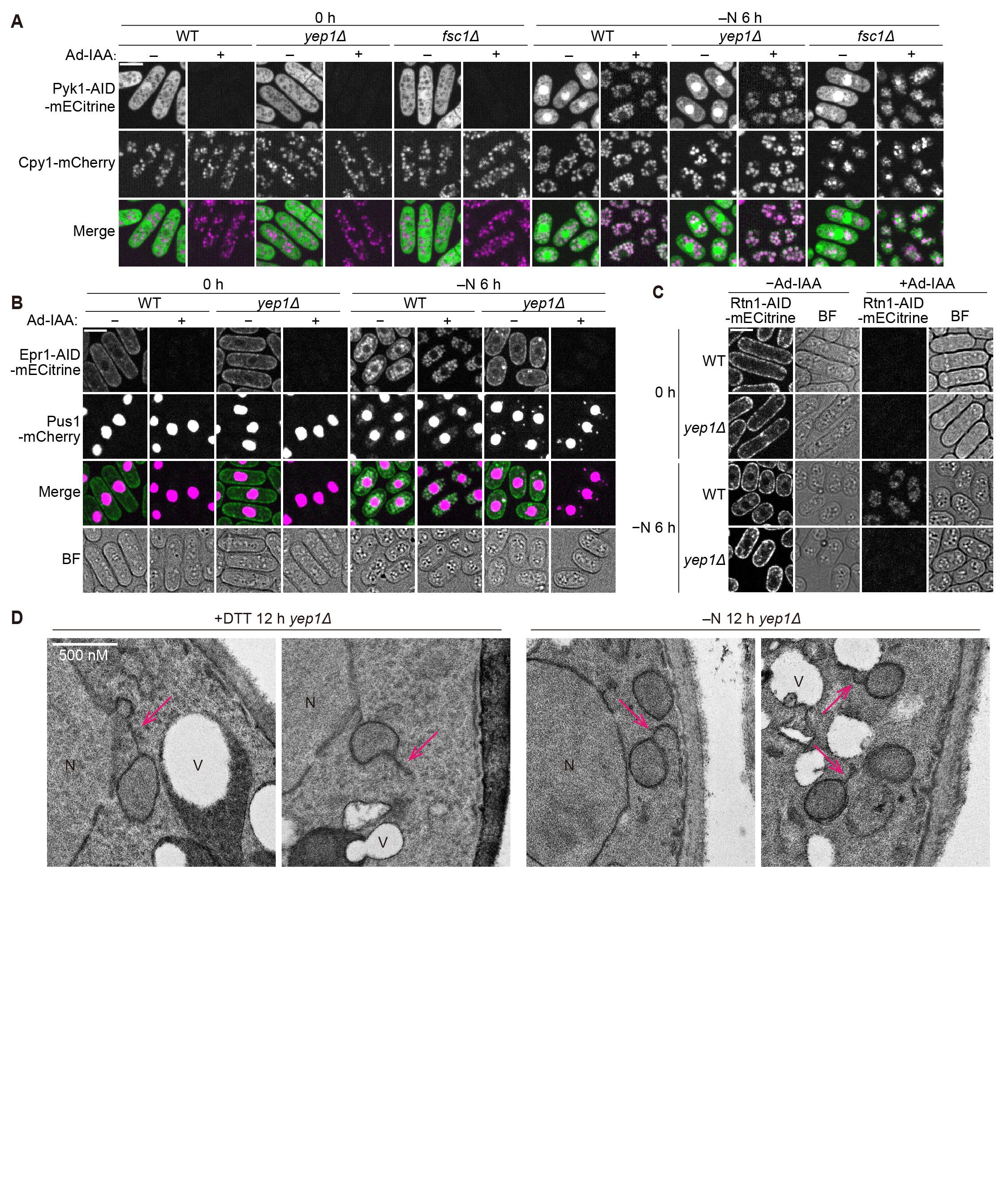

### Supplementary Figure S4

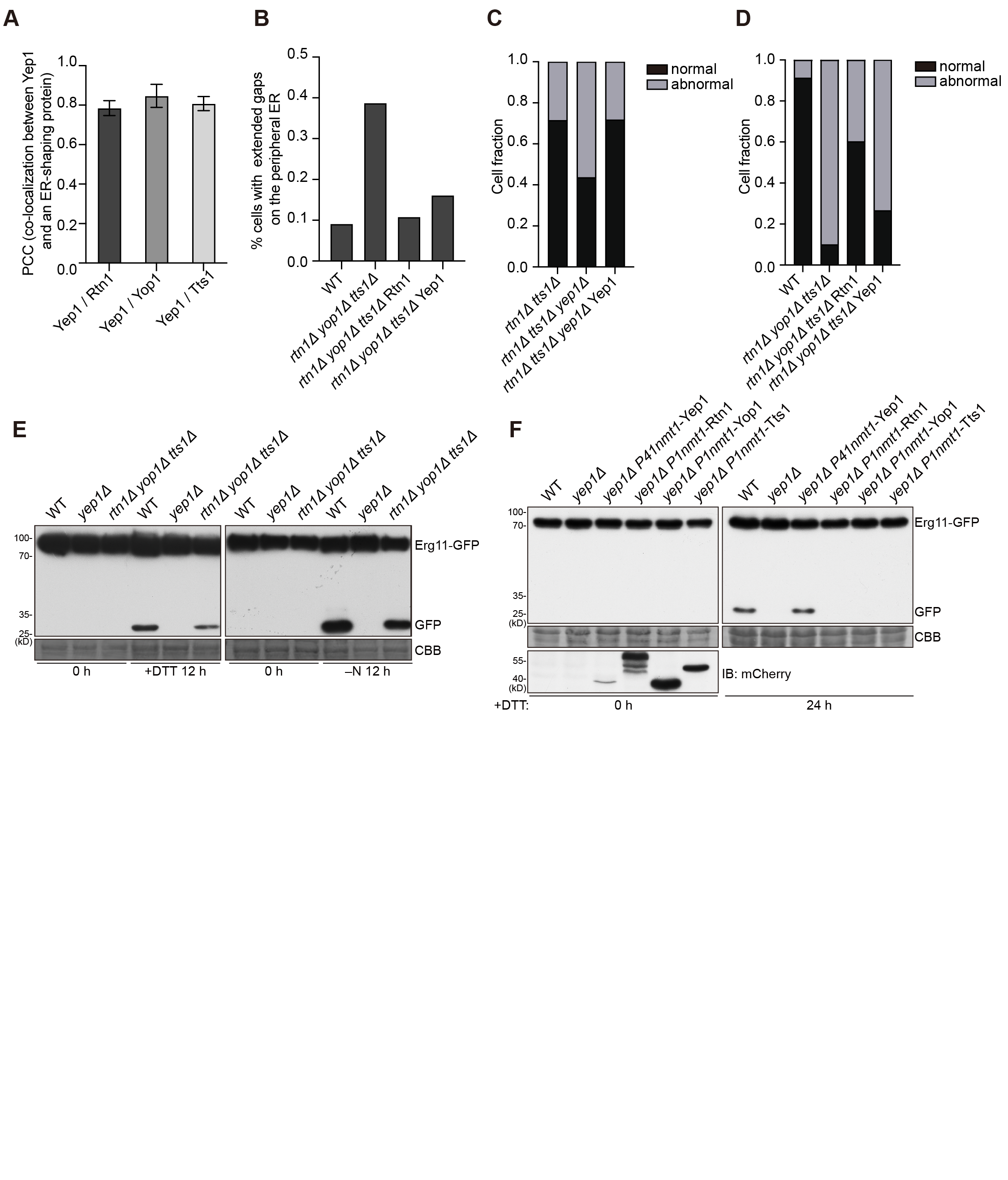

### Supplementary Figure S5

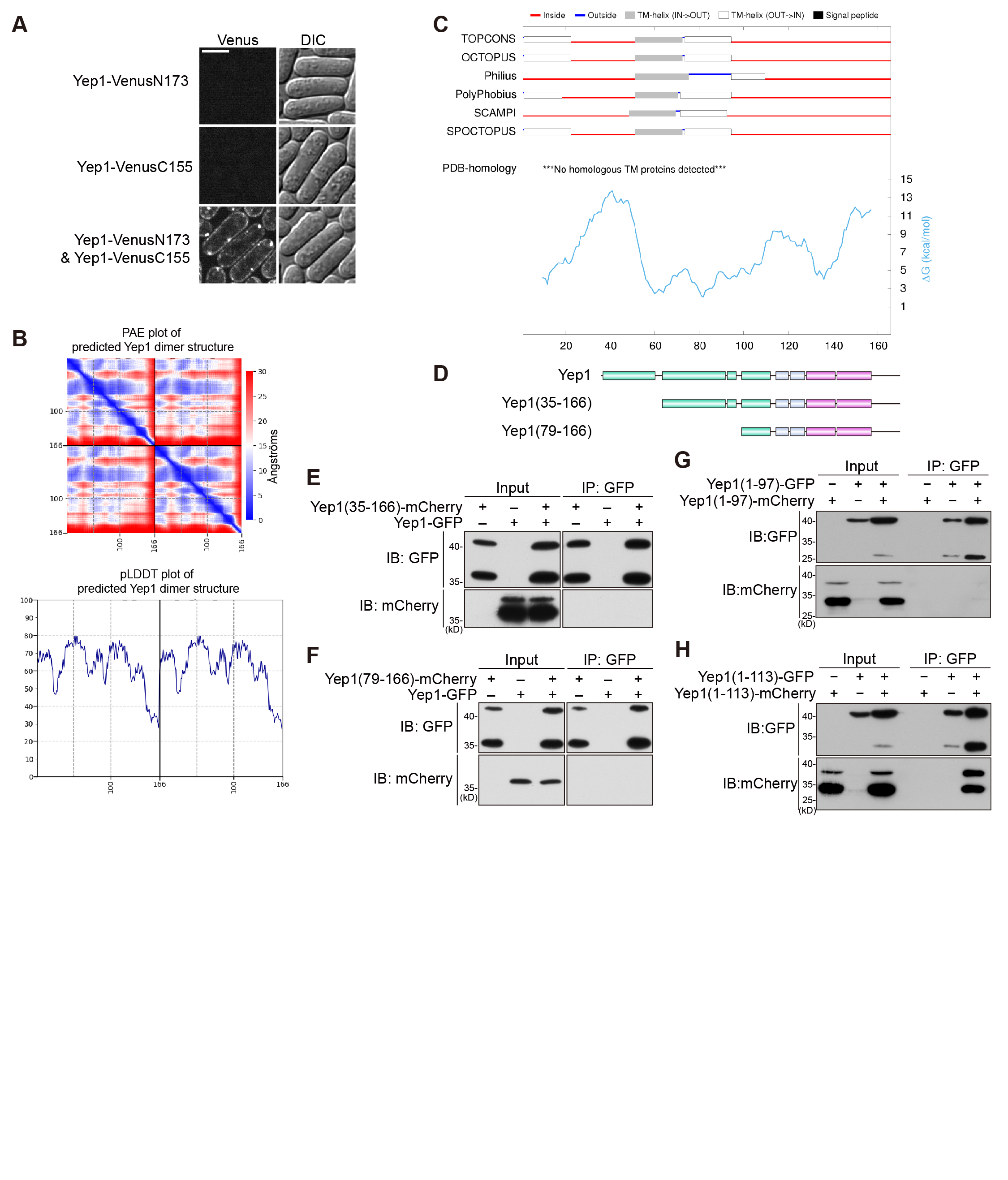

### Supplementary Figure S6

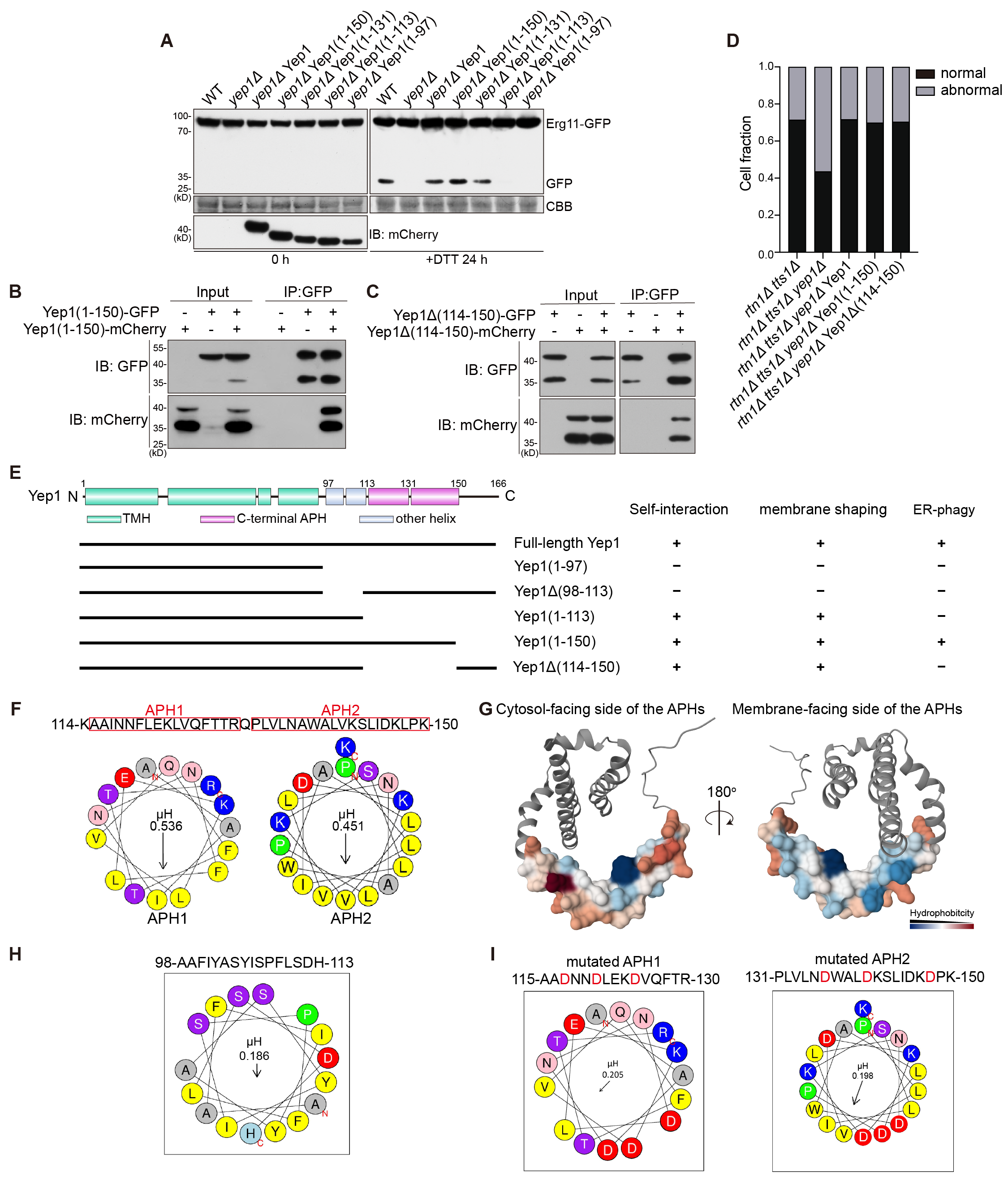

### Supplementary Figure S7

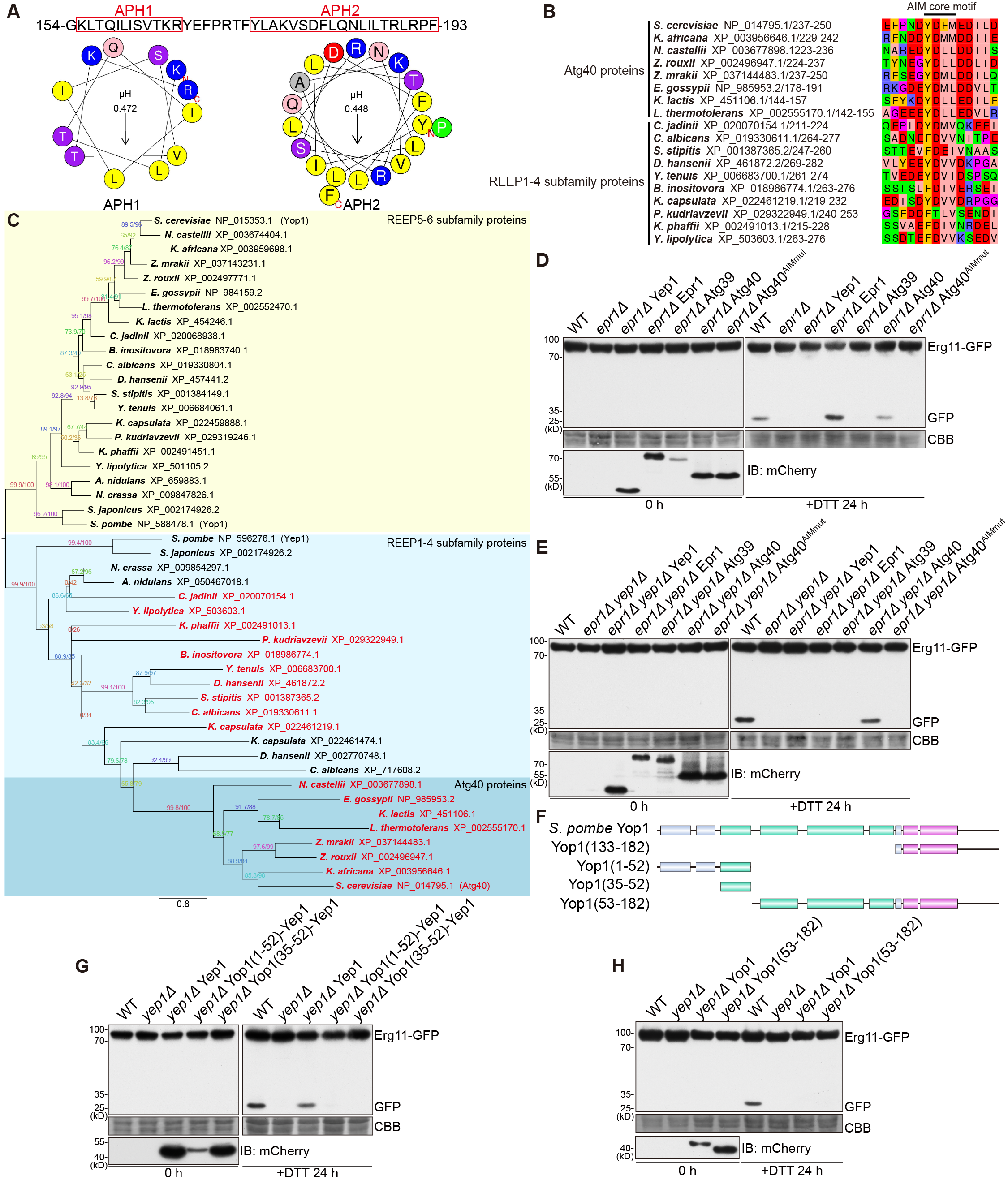

### Supplementary Figure S8

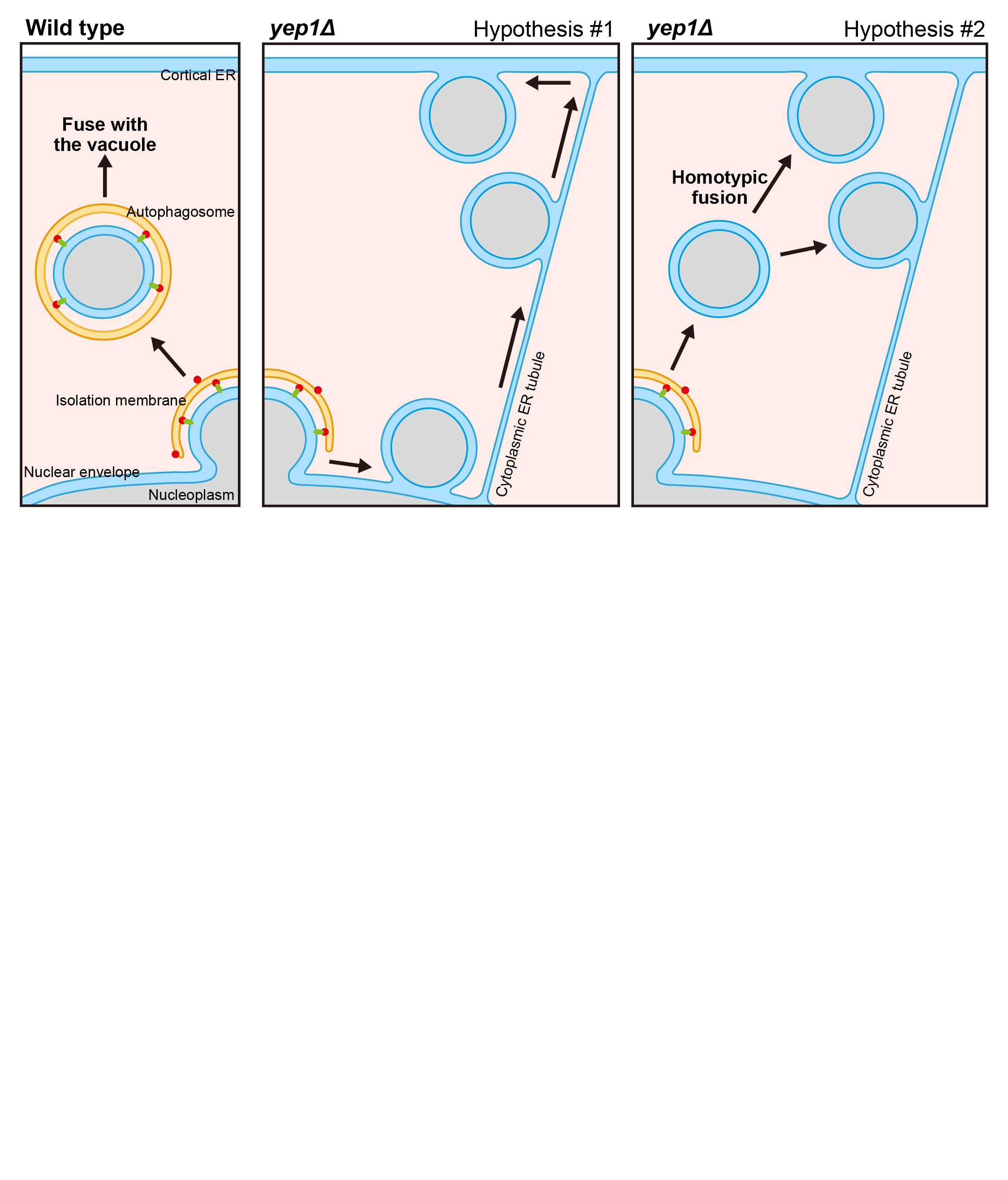
